# Mechanically Transduced Immunosorbent Assay To Measure Protein-Protein Interactions

**DOI:** 10.1101/2021.02.05.429999

**Authors:** Christopher J. Petell, Kathyrn Randene, Michael Pappas, Diego Sandoval, Brian D. Strahl, Joseph S. Harrison, Joshua P. Steimel

**Affiliations:** Department of Biochemistry and Biophysics, The University of North Carolina School of Medicine, Chapel Hill, NC 27599, USA; UNC Lineberger Comprehensive Cancer Center, 450 West Drive, University of North Carolina, Chapel Hill, NC, USA 27599; USA; Department of Chemistry, University of the Pacific, Stockton, CA 95211, USA; Department of Biological Engineering, University of the Pacific, Stockton, CA 95211, USA; Department of Mechanical Engineering, University of the Pacific, Stockton, CA 95211, USA

**Keywords:** METRIS, Rolling parameter, Ferromagnetic, Friction, K_*d*_, Protein-protein interactions, epigenetics, ubiquitin, UHRF1, DIDO1, ORC1

## Abstract

Measuring protein-protein interaction (PPI) affinities is fundamental to biochemistry. Yet, conventional methods rely upon the law of mass action and cannot measure many PPIs due to a scarcity of reagents and limitations in the measurable affinity ranges. Here we present a novel technique that leverages the fundamental concept of friction to produce a mechanical signal that indicates binding. The mechanically transduced immunosorbent (METRIS) assay utilizes rolling magnetic probes to measure PPI interaction affinities. METRIS measures the translational displacement of protein-coated particles on a protein-functionalized substrate. The translational displacement scales with the effective friction induced by a PPI, thus producing a mechanical signal when a binding event occurs. The METRIS assay uses as little as 20 pmols of reagents to measure a wide range of affinities while exhibiting a high resolution and sensitivity. Here we use METRIS to measure several PPIs that were previously inaccessible using traditional methods, providing new insights into epigenetic recognition.

## Introduction

Protein-protein interactions (PPIs) are essential to cellular biology and both high- and low-affinity interactions are required to maintain robust and dynamic responses in biological circuits [23, 29]. Low-affinity interactions are commonly leveraged, as is seen for multivalent recognition [27], readers of highly abundant proteins, and in protein allostery [8]. In particular, recognition of the epigenome is recognized to rely on the interplay between post-translational modifications (PTMs), like methylation, phosphorylation, and ubiquitination [28,32,51]. Furthermore, multidomain proteins are often regulated by allostery through weak interdomain interactions [18, 33]. Increasingly, the importance of weak interactions or relatively small changes in PPI affinity has been realized.

Despite the increasing sophistication of studying PPIs, biochemical characterization of these weaker and similar strength interactions remain a significant hurdle. Many techniques are useful for examining protein binding strength, each with its own set of limitations [35, 37, 45]. However, virtually all the commonly used techniques to measure biological interactions, like ELISA, FP, SPR, NMR, BLI, AUC, and ITC, rely on the law of mass action, and to measure protein binding affinities in the *µ*M range and above highly concentrated proteins or ligands are required [50]. For many systems obtaining such large quantities of materials can be unattainable and at these concentrations thermodynamic non-ideality occurs and proteins can aggregate, self-associate, and non-specific interactions occur, obfuscating the binding signal. [39, 48]. NMR is the gold standard method to measure weak interactions [26, 46], however in addition to requiring copious amounts of materials, the proteins must also be isotopically labeled, a single affinity measurement requires substantial instrument time and complex data analysis, and of all the methods mentioned is the lowest throughput. Another difficulty arising when measuring similar strength interactions, e.g., 3-5 fold differences. Several factors contribute to this limitation, but determining the active fraction of protein is significant because, for most fitting techniques, the calculated affinity is a dependent variable of the protein concentration [21, 22]. Another factor in differentiating similar strength interactions is that most binding measurements have low statistical power due to the resource intensiveness of performing multiple replicates. A method where binding strength can be measured independent of protein concentration and that has high statistical power would be valuable.

Here we present a novel approach to measuring the strength of biological interactions that is moderately high-throughput, requires a minimal amount of protein material, and can measure a wide range of K_*d*_ values from 10^*−*2^*−*10^*−*15^M. This technique was initially inspired by the rolling of biological cells, like neutrophils exhibiting haptotaxis on endothelial cells. Neutrophil motion is driven by chemical or ligand gradients [47]. The neutrophils roll on the endothelial cells due to PPIs between the cell surface receptors. The PPIs increase the effective friction between the two cells, allowing the rotational motion to be converted into translational displacement. We aimed to create a single particle biomimetic technique that leveraged this fundamental physical concept of friction to produce a mechanical signal to indicate binding events, the Mechanically Transduced Immunosorbent assay (METRIS). METRIS utilizes protein functionalized ferromagnetic particles to mimic the rolling cells. These ferromagnetic particles are made active via actuation of an externally applied rotating magnetic field and the particles proceed to roll, henceforth referred to as rollers, and translate across the surface using a similar mode of locomotion as the neutrophils. When the rollers are placed on a functionalized surface, the amount of rotational motion converted into translational motion depends on the effective friction between the rollers and the substrate. That effective friction scales with the strength of the binding interaction. Thus, a higher affinity PPI between the roller and the substrate will result in a larger translational displacement of the roller. Since both the roller and surface have immobilized proteins, the method is not dependent on mass action and requires approximately 20 pmols to measure PPIs regardless of their strength.

Using this METRIS assay, we reproduced well-characterized binding preferences for two different methyllysine histone reader domains [16, 24] and weak interactions between the E2 Ube2D [6] and UBL-domains [9]. These affinities range between 10^*−*4^*−*10^*−*6^M. However, we were also able to measure several weaker interactions between unmodified histone peptides, which allowed us to measure the Δ ΔGs for the phospho/methyl switch phenomenon in DIDO1-PHD [1]. Finally, we also show that this method can be used to measure a weak interdomain interaction between the isolated UHRF1-UBL domain and SRA domain, which is known to control the E3 ligase specificity and epigenetic DNA methylation inheritance [9, 13]. Collectively our results show that the METRIS assay can be a very powerful technique which has the potential to provide additional insight into PPI interactions that were not previously possible using other methods.

## Materials and Methods

### Magnetic Probe and Substrate Functionalization

The streptavidin coated ferromagnetic particles, provided by Spherotech, are composed of a core of polystyrene and CrO_2_. 10*µL* of the stock solution, 1.0% w/v, was extracted inserted into a micro-centrifuge tube, provided by Fisher Science. Biotinylated peptides were then inserted into the tube with the streptavidin coated ferromagnetic particles. The amount of peptides was such to coach each bead 50*×* the theoretical limit, 1mg of beads binds 0.16nmole of biotin, to ensure all binding sites on the beads were covered. The bead and peptide solution was left to react at room temperature for at least 2 hours.

The substrates are avidin coated glass slides, provided by Arrayit, with a ligand density of 1.1 *×*10^10^ ligands per mm^2^. Microfluidic channels were created on this substrate using two pieces of double sided tape, provided by 3M. The pieces of double sided tape were cut to a with of several mms and a length of at least 25mm. The pieces of tape were placed parallel to each other and at a distance of approximately 5mm apart. Then a glass coverslip, provided by VWR, was placed on top of the tape to create channels approximately 22*×* 5mm.

A solution of biotinylated proteins was then inserted into the channel. The amount of proteins inserted was again enough to coat the channel surface 50*×* the theoretical limit to ensure that all of the sites on the substrate were coated. The substrate and solution was left in a sealed container for two hours to allow the proteins time to bind to the substrate. After two hours the solution was washed from the channel to remove any excess protein that was not attached to the substrate. Then the solution of peptide coated ferromagnetic beads was diluted approximately 2000*×* to reduce the probability of two ferromagnetic beads forming a magnetic dimer that cannot be analyzed in the rolling parameter analysis. The channel was sealed with epoxy and magnetized by an external permanent neodymium magnet. The substrate was placed in the slide holder at the center of the Helmholtz Coil Inspired Experimental Apparatus.

### Helmholtz Coil Inspired Experimental Apparatus

Three pairs of coils were secured in an apparatus, made of aluminum T-slots, and attached to an optical breadboard. The coils have an inner diameter of 7cm and an outer diameter of 13cm, as seen in Fig. SS1.

### Protein purification and biotinylation

GST-[DIDO1-PHD/ORC1-BAH]-avi recombinant proteins we cloned into the pGEX-4T1 vector (GE, 27458001) to generate GST-[DIDO1-PHD/ORC1-BAH]-avi recombinant proteins. Recombinant proteins were purified as described in previous work [34]. Briefly, the recombinant proteins were induced to express in SoluBL21 cells (Fisher, C700200) after reaching an OD600 of 0.4 with 0.2 mM IPTG and by shifting to 16^*o*^C for overnight growth. After induction, the cells were pelleted and resuspended in Lysis Buffer (50mM HEPES, 150mM NaCl, 1mM DTT, 10% glycerol, pH 7.5) supplemented with protease inhibitors, then incubated in the presence of lysozyme (Sigma, L6876) and nuclease (ThermoFisher, PI88700) for 30 minutes. After this the cells were sonicated for six rounds consisting of 10 seconds continuous sonication at 50% intensity, 50% duty cycle followed by 60 seconds on ice. Lysates were centrifuged for 10 minutes at 10,000 rpm and the clarified lysates loaded onto a glutathione resin and purified by batch purification according to the manufacturer’s protocol (ThermoFisher, PI16101). Purified proteins were then dialyzed against Lysis Buffer to remove GSH and quantified using a Bradford assay per the manufacturer instructions (BioRad, 5000006) prior to being stored at −80^*o*^C. Ube2D1 is a his-tagged protein that was purified according to previous publications through standard Ni-NTA purification. The UHRF1-UBL, W2V-mutant, and UHRF1-SRA domain were cloned into a modified version of His-MBP-pQE80L vector that we have previously described. For the UHRF1-UBL domain and W2V mutant an N-terminal cystine was added using PCR for chemical conjugation with maleamide. These proteins were grown to O.D. 0.6 and induced with 0.6mM IPTG. MBP was cleaved using TEV purified in house and removed using anion exchange. The ubiquitin with an N-terminal cystine was purified using a pGEX-4T1 expression system described here. The ubiquitin was removed from the resin by cleavage with TEV. Purified proteins with an avi-tag were biotinylated by using BirA following the BirA500 kit’s protocol (Avidity, BirA500). Biotinylation was confirmed by performing a Coomassie gel shift assay according to Fairhead and Howarth, 2015 [12]. Cysteine Biotinylation was carried out using Poly(ethylene glycol) [N-(2-maleimidoethyl)carbamoyl]methyl ether 2-(biotinylamino)ethane (Sigma 757748) (Biotin-maleamide). Typically small volumes were biotinylated such that very little biotin-malamide was needed (below a mg) so we added some powder and confirmed biotinaylation with SDS-page gel. For UHRF1-UBL variants and ubiquitin there is only a single engineered cysteine available for modification. For the Ube2D, UHRF1-SRA domain, and GST-PHD-DIDO1 we labeled native cysteines which resulted in heterogenous labeling. Excess biotin-maleamide was removed using size-exclusion or anion exchange for the UHRF1-UBL and ubiquitin, and dialysis for the SRA and GST-PHD-DIDO1. Proteins were typically aliquoted and frozen before METRIS analysis. Both methods were evaluated for their ability to return RP values within error, which is shown in Fig. SS3.

### Histone peptide microarrays

Histone peptide microarrays were performed and analyzed as described in Petell et al., 2019 [34]. In brief, 500 nM of the avi- and GST-tagged DIDO1-PHD or ORC1-BAH constructs in 1% milk 1x PBST (10 mM Na2HPO4, 1.8 mM KH2PO4, 2.7 mM KCl, 137 mM NaCl, pH 7.6, 0.1% Tween-20) were incubated overnight at 4^*o*^C with shaking. The following day, the arrays were washed by submerging in 1x PBS briefly, then submerged in 0.1% formaldehyde in 1x PBS for 15 seconds to cross-link, formaldehyde was then quenched by submerging in 1 M glycine in 1x PBS for one minute, after which the arrays were submerged in 1x PBS and inverted five times to remove remaining glycine. Next, the arrays were washed three times with high-salt 1 X PBS (1x PBS with 497 mM NaCl rather than 137 mM NaCl) for 5 minutes each at 4^*o*^C with shaking. Then, the arrays were incubated with a 1:1000 dilution of anti-GST (EpiCypher, 13-0022) in 1% milk 1x PBST for two hours at 4^*o*^C with shaking. After incubation with anti-GST antibody the arrays were washed with 1x PBS, three times for five minutes at 4^*o*^C with shaking. Next, they were exposed to a 1:10,000 dilution of anti-Rabbit AlexaFluor-647 (Invitrogen, A21244) for 30 minutes at 4^*o*^C with shaking. Lastly, the arrays were washed three times for five minutes with 1x PBS as in the previous wash step, then submerged in 0.1x PBS prior to imaging. The arrays were imaged using a Typhoon (GE) and quantification was carried out using ImageQuant TL software. Analysis of the data was done by first averaging the triplicate intensities for a given peptide on the array; the values for an arrays’ dataset were then linearly scaled from 0 to 1 by applying a min-max formula such that the minimum value became 0 and the maximum 1. After, this all the scaled array values were combined to derive a single average and standard deviation for each peptide and the averages used for the graphs; see plots for what peptide modification states are shown. For the average and standard deviations of each individual peptide, see the Supplemental Data File. Results for the DIDO1-PHD and ORC1-BAH domains showing all peptides carrying the specified modifications, alone and in combination with other PTMs is shown in Fig. SS2.

### Data Collection and Statistical Analysis

All experiments for METRIS were performed at least 12 times in replicate and all array data consist of at least 3 replicates, and averages with standard deviation are shown in the tables for each figure. All statistical analysis were done by using the Student’s T-Test (unpaired, two-tailed distribution). The results of this statistical analysis are reported in Fig S.S4.

## Results

### Rolling Parameter Scales with Interaction Affinity of PPI

In the METRIS assay, rollers are placed in a Helmholtz coil inspired apparatus (see Fig. 1A and S1) where an externally rotating magnetic field is applied at a constant frequency, *ω*. The permanent magnetic moment of the roller couples with the applied magnetic field, producing a magnetic torque and subsequent rotation of the ferromagnetic bead [42, 44]. Without friction, the rollers would rotate mostly in place with the frequency of the applied magnetic field; however, friction between the rollers and the substrate will convert some of that rotational motion into translational displacement, Δ*x*, thus indirectly measuring the friction between the substrate and the rollers. In this system, friction is determined by the strength and density of PPIs between the roller and the coated substrate. Thus, the translational displacement will scale with the density and affinity of the PPIs being measured. However, the displacement is also a function of the diameter of the roller, *D*, and the frequency of rotation of the applied magnetic field, *ω*. Here we define a dimensionless parameter to account for these parameters that we refer to as the rolling parameter, RP, *ζ*, as seen in Fig.1B

**Figure 1.**
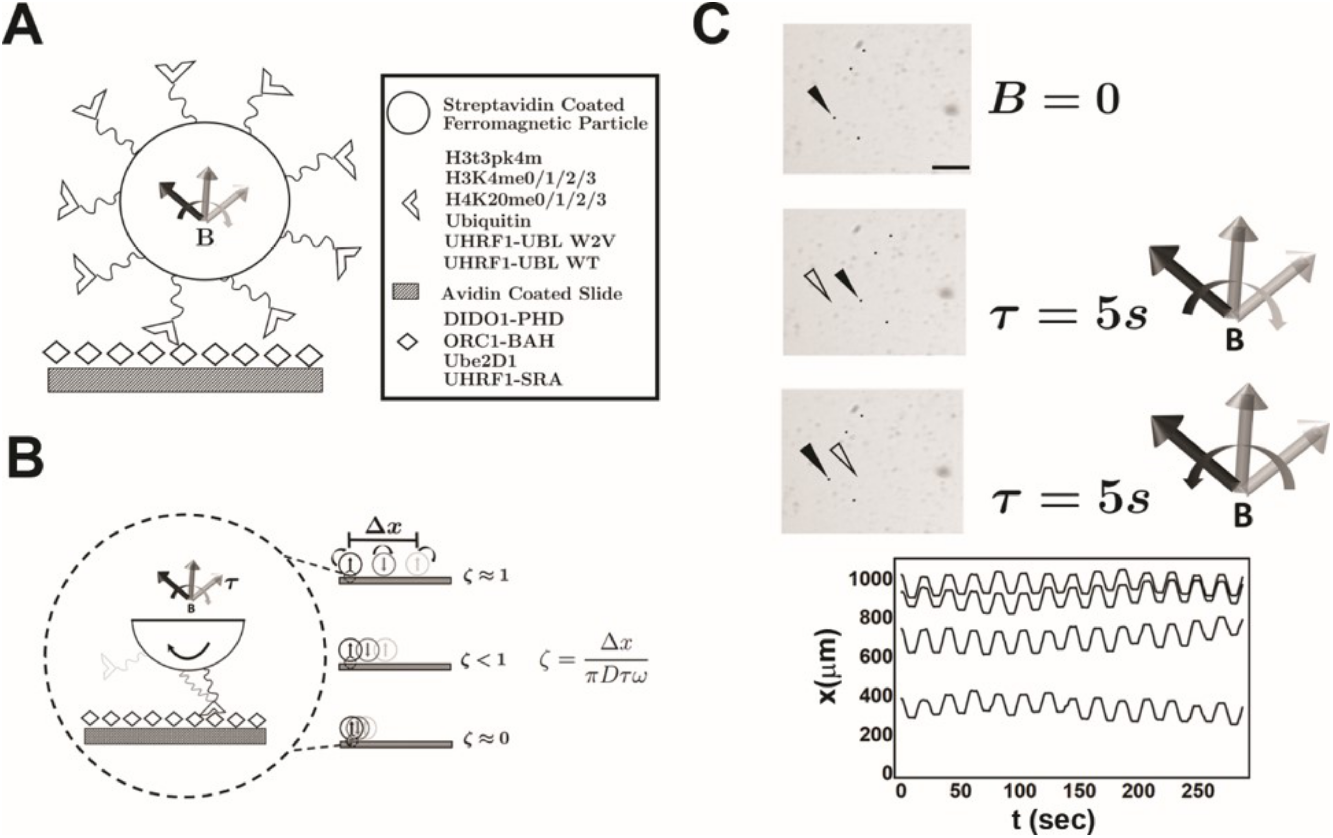
Experimental schematic of Mechanically Transduced Immunosorbent Assay (METRIS) assay used to measure protein-protein interactions.. A) Diagram of the substrate and ferromagnetic functionalization protocol. B) Schematic of the translational displacement of rollers due to effective friction between the probes and substrate, which scales with interaction affinity. The translational displacement is then analyzed to calculate a rolling parameter (RP), *ζ*, that is used to measure binding affinity. C) Representative microscopy images (scale bar in black is 100*µ*m) of rollers (black points) prior to magnetic field actuation, (top), after actuation in clockwise (middle), and after actuation counter-clockwise (bottom). The graph summarizes the roller trajectories for the duration of the experiment.

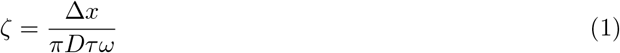

where Δ*x* is the translational displacement of the roller, *D* is the diameter of the roller, *τ* is the actuation period of the magnetic field, and *ω* is the rotational frequency of the magnetic field. The density of the interactions between the roller and the substrate are kept as constant as possible from experiment to experiment by fully saturating both the rollers and the substrate with proteins and peptides. As described in the SI, both the rollers and substrate are coated 50*×* the theoretical number of binding sites, so virtually all of the sites should be occupied. Additionally, a series of washing steps are carried out to make sure no unbound protein or peptide remains on the surface. If the surface was not uniformly functionalized, the roller’s displacement in these regions would be detected by correlations to either the individual roller or area on the substrate. However, no such anomalies were observed in these experiments.

To measure the RP of the rollers, a clockwise field was actuated at *ω* = 1*Hz* for *τ* = 5 seconds. The field was then turned off for *τ* = 5 seconds. A counter-clockwise field was actuated at *ω* = 1*Hz* for *τ* = 5 seconds and then the field was turned off for *τ* = 5 seconds again. This process was repeated 18 times, and several example images of rollers and roller trajectories can be seen in Fig.1C. The rolling parameter is calculated from the observed roller displacement divided by the maximum theoretical velocity of a rolling sphere where all the rotational torque is converted into translation, so the rolling parameter varies from 0-1. A rolling parameter of 0 corresponds to a surface with no effective friction. Experimentally, a rolling parameter of 0 is never observed due to hydrodynamic friction between the roller and the substrate. We approximate the rolling parameter (0.081 *±* 0.004) observed for a system consisting of a streptavidin coated roller rolling on an aviding coated substrate in 1*×* PBS to be to be the null interaction scenario. While it is impossible to know the true K_*d*_ value for a null interaction, the weakest PPI measured are in the 10^*−*2^M range [52] and enzymes with K_*d*_ values in the 10^0^M range have been reported [2] so we can assume that null interaction must be between 10^0^M and the concentration of water 5.5*×* 10^2^M, and we settled on 10^0^M as an estimation of the null interaction. A rolling parameter of 1 corresponds to a binding affinity that is extremely large with high effective friction, our best approximation of this is the interaction between biotin and streptavidin, K_*d*_=10^*−*15^M, for which we observe a RP of 0.918 *±* 0.002.

### DIDO1-PHD Phospho/Methyl Switch Characterized by METRIS

Previous studies have demonstrated the utility of the METRIS assay to measure rolling parameters for a variety of interactions (e.g., Protein A-Fc, histidine-metal, and protein-PIP lipid interactions) [42–44]. Here, we wanted to see how well we could reproduce binding strengths for known PPIs and determine the robustness of the METRIS assay. We focused our attention on weak interactions and interactions between several protein pairs that are similar in binding strength. We first examined the well-established interaction between DIDO1-PHD and H3K4 methylation. DIDO1 is responsible for interchanging between active and silent chromatin states in embryonic stem cells, and its chromatin localization is regulated through a phospho/methyl switch, where phosphorylation of H3T3 evicts DIDO1 from chromatin during mitosis [11, 14, 25]. The affinities for mono-, di-, and trimethylated peptides are well described in the literature [16] and interactions with the unmodified peptide and H3T3pK4me3 were too weak to be measured in the experiment setup. H3K4 peptides and DIDO1-PHD were both immobilized to the rollers and substrate through biotin-streptavidin interactions. The H3 N-terminus (a.a. 1-20) was biotinylated and coated on the roller, and biotinylated avi-tagged GST-DIDO1-PHD was attached to the substrate (see SI Methods for details). DIDO1 has a preference for H3K4me3 *<* H3K4me2 *<* H3K4me1 [16]. The measured rolling parameters match this preference, with the largest rolling parameter for K4me3 (0.233*±*0.012) *<*H3me2 (0.213*±* 0.010) *<* H3me1 (0.176*±*0.005) and H3 and H3T3pK4me3 being the lowest, although still above the baseline rolling parameter value of 0.081 as seen in Fig.2A. While the overall change to the RPs is small, these differences are all statistically significant because the data set has good statistical power and small percentage errors (*<*5%) (Supplemental Fig.4A). In order to correlate binding affinity to RP, a log-log plot of K_*d*_ vs. RP showed a linear relationship between the three known DIDO1-PHD binding interactions to the methylated peptides (R^2^=0.995). This fitting also included a no-binding avidin-streptavidin interaction (RP=0.081) estimated to have a K_*d*_ = 1M and the streptavidin-biotin interaction where K_*d*_ = 10^*−*15^M [10] (Fig.2B). Strikingly, this experiment shows a linear dependence of the log of the RP to the log of K_*d*_ over roughly fifteen-orders of magnitude.

**Figure 2.**
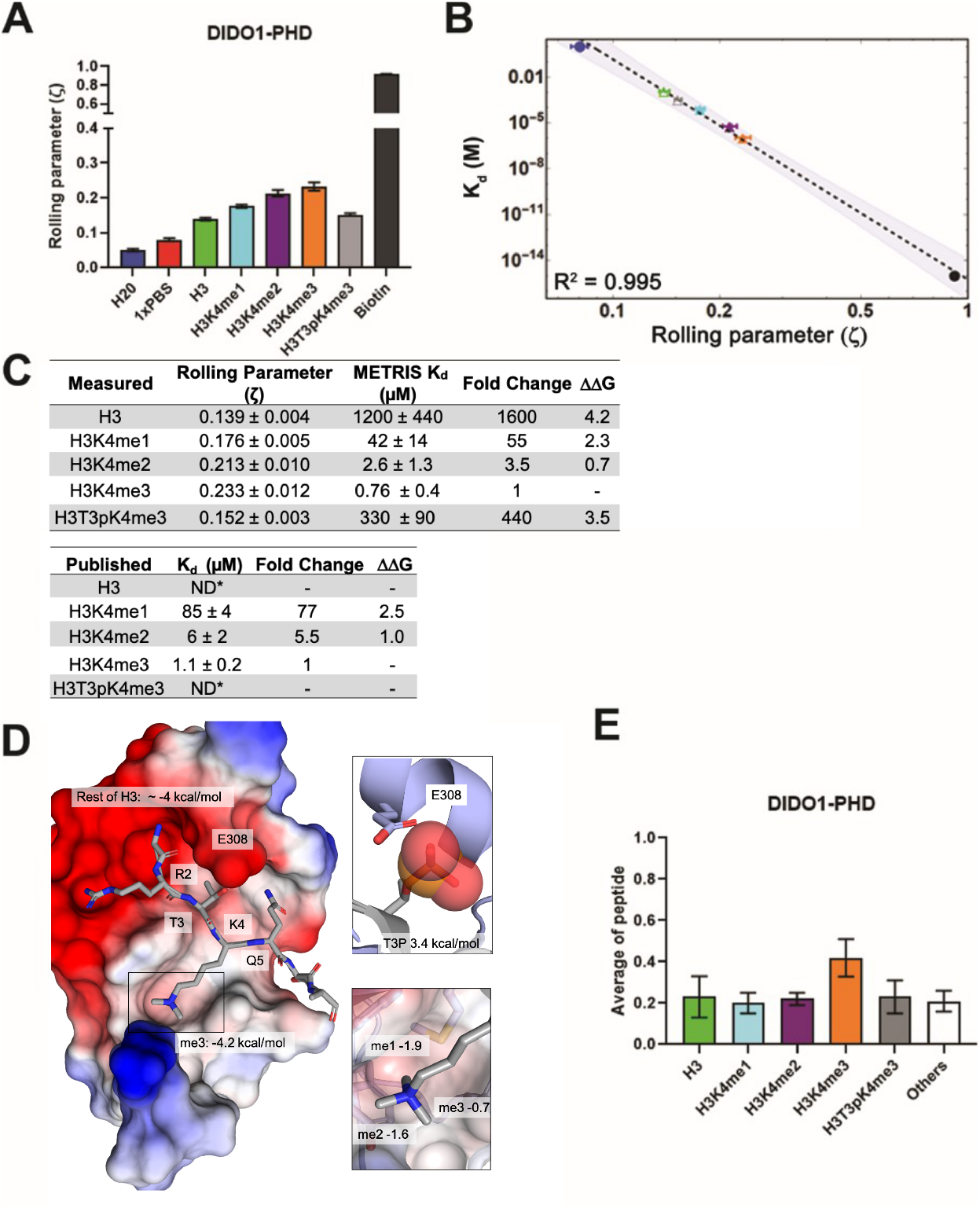
DIDO1-PHD interactions with H3 peptides characterized using METRIS A) Results of METRIS experiments using the DIDO1-PHD and the indicated H3K4 methylated peptide or controls. See Figure S4A for results of statistical analysis; all comparisons are significant. B) Log-Log plot of the rolling parameters, *ζ*, from panel A. Extrapolated point markers are unfilled, and the 95% confident interval for the fitting is depicted. C) Table of rolling parameters and associated K_*d*_ estimates for the DIDO1-PHD interactions. Fold change is calculated as the ratio between the K_*d*_ values for the indicated peptide and for H3K4me3. These ratios are used to calculate ΔΔ*G* at T=298K. The published values are from Gatchalian et al., 2013 using NMR (me1) and tryptophan fluorescence (me2/3); *ND= Not determined. D) Image of the DIDO1-PHD crystal structure with H3K4me3 peptide, with the PHD surface electrostatic potentials shown (red = negative, blue = positive), the ΔΔ*G* for K4me3, and the estimated ΔΔ*G* for the rest of the peptide. The PTM reader sites are shown with greater detail to the right. Here ΔΔ*G* is calculated between the sequential methyl states, and the ratio of H3T3pK4me3 and H3K4me3 give the ΔΔ*G* for T3p. E) Results of the peptides from panel A are shown from a histone peptide microarray assay using DIDO1-PHD (see S3A for complete peptide plot). Only H3K4me3 is statistically significant (see Figure S4A for results of statistical analysis). While these results indicate general binding trends, they cannot provide K_*d*_ estimates and do not have high enough resolution to distinguish between weaker binding interactions.

There is a clear correlation between RP and the measured K_*d*_, the equilibrium constant for interactions, despite METRIS being a non-equilibrium technique. K_*d*_ is a ratio between the first-order dissociation rate (K_*off*_) and the second-order association rate (K_*on*_) [40]. For most PPI, the K_*on*_ rates are very similar, and thus the K_*d*_ constant is mostly dependent on K_*off*_. However, kinetic constants for binding interactions are rarely reported since few techniques can access this information, so for many interactions, only K_*d*_ is known. Since we do not have a theoretical model that relates RP to K_*d*_, we sought to use an empirical fitting method based on the excellent correlation we observed between RP and K_*d*_ (Fig.2B). *Using this fitting method, we could accurately reproduce the literature K*_*d*_ values with high accuracy; all of the predicted K_*d*_ values were roughly 2-fold tighter than the established NMR values [16] and the fold difference between different methylation states similar (Fig.2C). Remarkably, we were also able to estimate METRIS-K_*d*_ values for the weak interaction between the H3T3pK4me3 peptide (340*µ*M *±* 90) and the unmodified H3 tail (1200*µ*M *±* 440). While these are empirically derived estimates for K_*d*_, it is clear from the RP measurements that these interactions are statistically distinct, and they represent a missing piece of data that is critical to a quantitative understanding of epigenetic recognition. The utility of this data is exemplified when evaluating the ΔΔ*G*^*o*^ (Δ ΔG) values, a common way to report the energetic contributions of individual amino acids for a set of related PPIs. Δ ΔG is calculated by taking the natural log of the ratio of two K_*d*_ values (K_*d*1_ and K_*d*2_ in equation 2) in the Gibbs free energy equation, where R is the gas constant and T is the temperature in Kelvin [41].

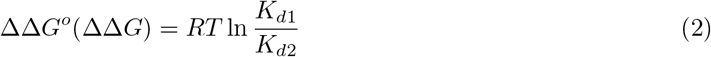

This analysis allows for calculating the energetic contributions of the individual PTMs. For example, K4me3 is worth 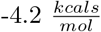 while T3P is worth 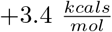 (Fig.2C). *To our knowledge, this is the first* energetic analysis of the DIDO1 phospho/methyl switch. These values have more context when viewed with the crystal structure of DIDO1-PHD (Fig.2D) [16]. The hydrophobic trimethyl-lysine binding site accounts for a significant amount of the total binding to the peptide, however, there are clearly other residues on H3 that interact with DIDO1-PHD, such as the N-terminus, R2, and T3, and therefore, it is not surprising that unmodified H3 can still bind and account for roughly 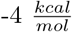 when using 1M K_*d*_ as the null reference. The deleterious effect of T3p is also resolved, since residue E308 of the PHD domain would clash and repel a T3p modified histone tail. Furthermore, this analysis also provides new insights into discrimination of methylation states by the DIDO1-PHD. For example, the greatest change in Δ ΔG occurs between H3 from H3K4me1 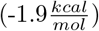, then H3K4me1 versus H3K4me2 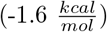, and H3K4me2 from H4me3 is the weakest 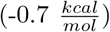. Thus, despite the DIDO1-PHD having the highest affinity for H3K4me3 it has the greatest discrimination between non-methylated H3K4 versus H3K4me1. The structure agrees with this observation, where two of the methyl binding sites are the most buried and the third is the most exposed one.

One of the significant advantages of the METRIS assay is that only 10 *µ*l of 2 *µ*M (20 pmol) is required to load the substrate and less is needed for the rollers. We compared METRIS to histone peptide microarrays, which is another methodology that can produce binding data with a minimal amount of protein (e.g., 500 *µ*l of 0.5 *µ*M (250 pmol) protein). While microarrays offer high-throughput screening, they lack the sensitivity to determine weak binding and small affinity differences. For DIDO1-PHD, we could observe a statistically significant difference between H3K4me3 and the other methylation states, but there were no other statistically significant differences (Fig.2E, S3A, and S4A). Given this result, METRIS is significantly more sensitive and quantitative than other common methods to measure protein affinities that use comparable amounts of reagents at low concentrations.

### Determining ORC1-BAH Methyl Preferences Using METRIS Analysis

We further validated the METRIS assay against using another methyllysine reader, the BAH domain of ORC1. ORC1 functions in licensing origins of replication by discriminating H4K20me2 from H4K20me1, a PTM on active chromatin, and H4K20me3 a repressive PTM [3, 4, 24]. We selected ORC1 because the reported affinities are within an order of magnitude, with a 2-fold difference reported between H4K20me1 and H4K20me3. The RP values we obtained matched the published binding preferences [24] H4K20me2 (0.263*±*0.011) *<* H4K20me1 (0.226*±*0.008) *<*H4K20me3 (0.215 *±*0.005)*<* H4 (0.202*±*0.005) (Fig.3A). Using the same fitting method, we observe a linear log-log dependence (R^2^ = 0.967) and the METRIS calculated K_*d*_ values were between 4-8 fold tighter than the published values, yet there was good agreement between the fold-change and accordingly the Δ ΔGs. (Fig.3B and C). Thus, the METRIS assay is sensitive enough to measure changes that are 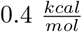.

**Figure 3.**
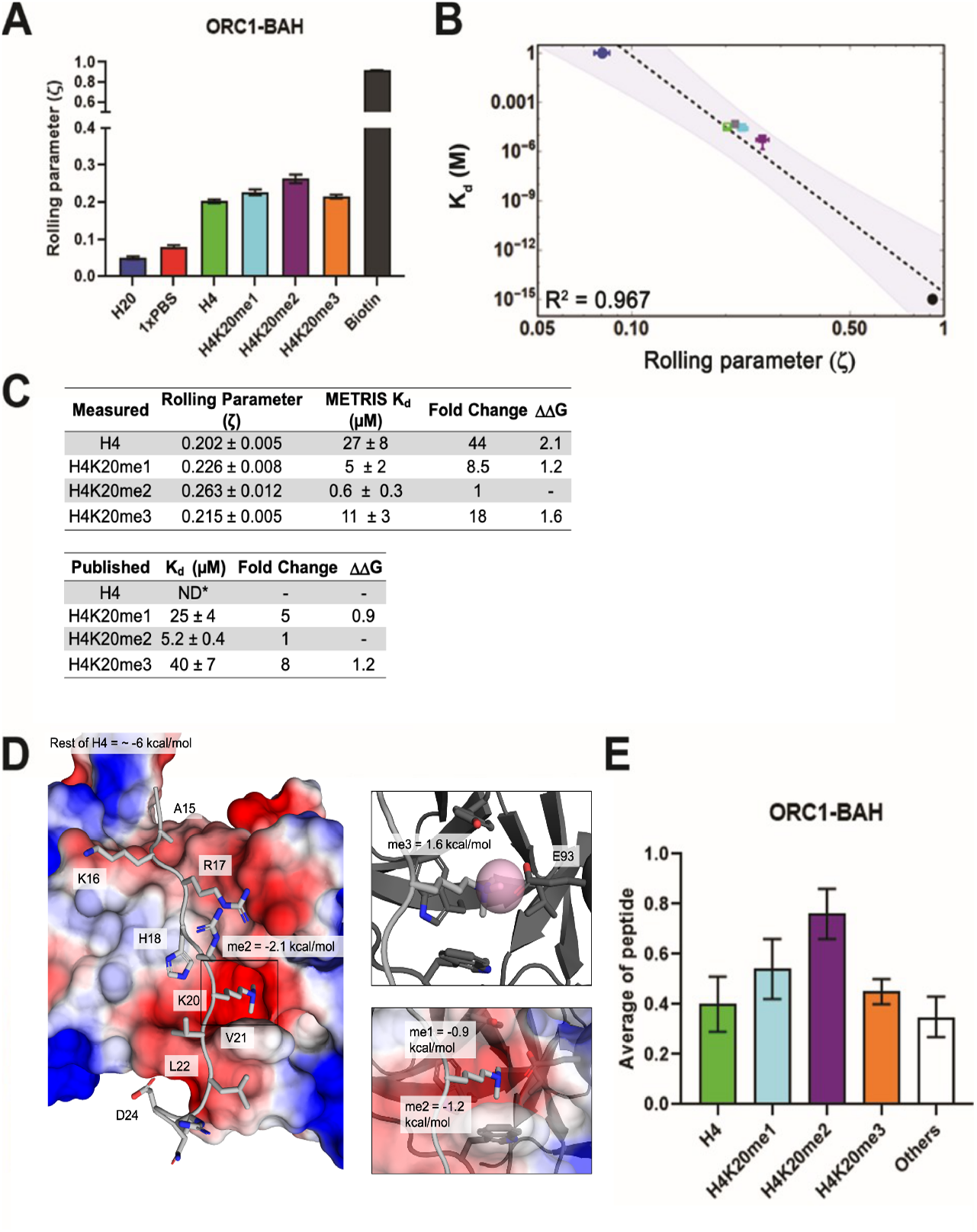
ORC1-BAH domain interactions characterized using METRIS. A) Results of METRIS experiments using the ORC1-BAH and the indicated H4K20 methylated peptide or controls. See Figure S4B for results of statistical analysis; all comparisons are significant. B) Log-Log plot of the rolling parameters, *ζ*, from panel A. Extrapolated point markers are unfilled and the 95% confident interval for the fitting is depicted. C) Table of rolling parameters and associated K_*d*_ estimates for the ORC1-BAH. Fold change is calculated as the ratio between the K_*d*_ for the indicated peptide and the K_*d*_ for H4K20me2. These ratios are used to calculate ΔΔ*G* at T=298K. The published values are from Kuo et al., 2012 using ITC; *ND= Not determined. D) Image of the ORC1-BAH crystal structure with H4K20me2 peptide, with the BAH surface electrostatic potentials shown (red = negative, blue = positive) as well as the ΔΔ*G* for K20me2 and the estimate for the rest of the peptide. The PTM reader site is shown with greater detail to the right. Here the ΔΔ*G* is calculated between the sequential methyl states. E) Results of the peptides from panel A from histone peptide microarray assay using ORC1-BAH (see S3A for complete peptide plot). Only H4K20me2 is statistically significantly different from the other H4 peptides. Again, we see that microarrays can indicate general binding trends but they cannot provide K_*d*_ estimates and do not have high enough resolution to distinguish between weaker binding interactions.

Using the METRIS assay, we could also measure binding to the unmodified H4, which has previously not been detected, and we measured it is 44-fold weaker than H4K20me2. With this value we could calculate that the Δ ΔG for K20me2 is worth 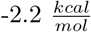. When comparing this to the DIDO1-PHD, we find that DIDO1-PHD has a stronger interaction with the PTM (−4.2 versus 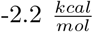) however the ORC1-BAH domain has a stronger interaction with the unmodified histone than the DIDO1-PHD (−6 versus 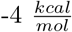). Examining the structure of ORC1-BAH domain bound to H4K20me2 [24] shows the methyllysine binding pocket is more charged than DIDO1-PHD, and likely, in part, contributes to the higher affinity to the unmodified peptide (Fig.3D). The METRIS analysis also furthers our understanding of ORC1-BAH discrimination amongst methyl states. We find the greatest differentiation between H4K20me2 and H4K20me3 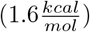 consistent with the biological role of ORC1 and this methyl sensing occurs through residue E93 (Fig.3D).

We also performed histone peptide microarrays on ORC1-BAH for comparison against the METRIS assay. The only statistically significant difference is between H4K20me2 and the other peptides (Fig.3E and S3 B), although the trends do match the literature and METRIS values, including the signal for the unmodified peptide when compared to the other peptides on the array, which support our findings with METRIS. However, due to the large standard deviation observed on the microarray, the assay would need to be repeated multiple times to achieve statistical significance. This highlights another advantage of METRIS assay, since it is a single particle method and the RP measurements are taken 38 times for each particle, this method has high statistical power.

### Investigating Noncovalent Interactions between Ubiquitin-like Domains and Ube2D1 Utilizing METRIS

We next used METRIS to investigate interactions with the protein post-translational modification ubiquitin. Ubiquitin has an expansive cellular regulatory role that is controlled by weaker interactions with effectors [7, 30] including non-covalent interactions with E2s and E3 ligases [5, 53]. Ubiquitin binding is wide-spread, and there are hundreds of UBLs in the human genome for these readers to discriminate amongst [20]. For example, the E2 Ube2D1 binds to ubiquitin noncovalently with an affinity of 206 *±* 6 and we have shown that a ubiquitin-like domain (UBL) on the E3 UHRF1 can bind with higher affinity (15 *±* 1 *µ*M with NMR or 29.0*µ*M *±* 1 with ITC) [9]. To probe this interaction with METRIS, both ubiquitin and the UHRF1-UBL domain were labeled using biotin-PEG-maleimide at an N-terminal cysteine installed for labeling, and Ube2D1 was labeled at native cysteines. Our RP data match the affinity trend UHRF1-UBL (0.131 *±*0.005) *>* ubiquitin (0.108*±* 0.004) (Fig.4A) and fitting METRIS-K_*d*_s produced values that were 5-fold weaker than the published values, but were in exact agreement with the 13-fold difference 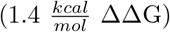 reported in the literature (Fig.4B, and S4C). Therefore we have demonstrated that METRIS can measure and distinguish interactions in the 10^*−*4^M range without utilizing highly concentrated protein solutions, providing a simple method to measure weak interactions.

**Figure 4.**
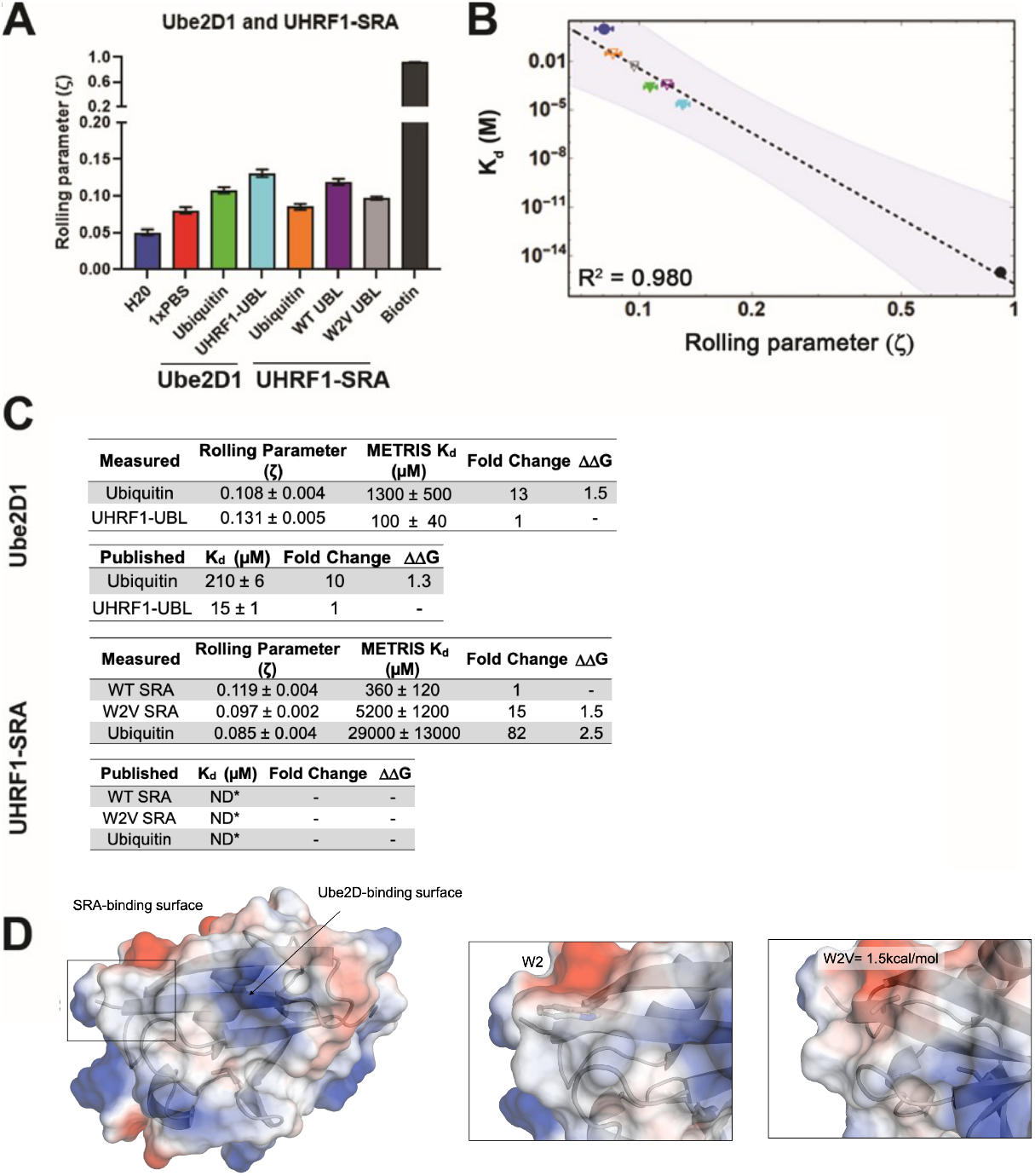
Measuring the interaction of UBL domains with Ube2D1 or the UHRF1-SRA domain using METRIS. A) Results of METRIS experiments measuring ubiquitin and the UHRF1-UBL domain binding to UbeD21 (E2) or the UHRF1-SRA domain. All comparisons are statistically significant (See Figure S4C for results of statistical analysis.) B) Log-Log plot of the rolling parameters, *ζ*, from panel A. Extrapolated point markers are unfilled and the 95% confident interval for the fitting is depicted. C) Table of all rolling parameters and associated METRIS-K_*d*_ estimates. Fold change is calculated as the ratio between the indicated protein and the UHRF1-UBL domain and the ΔΔ*G* is calculated using these ratios at T=298K. K_*d*_ values for ubiquitin are taken from Buetow et al., 2015 and UHRF1-UBL value taken from DaRosa et al., 2018. D) Image of the UHRF1-UBL binding surface for the UHRF1-SRA and Ube2D1. shown with electrostatic surface potentials (red = negative, blue = positive) with insets highlighting the change of the UBL surface with the W2V mutation and the associated ΔΔ*G*.

### Direct Measurement of an Interdomain Interaction Between UBL and SRA Domains of UHRF1 using METRIS

For epigenetic readers/writers, there is an abundance of examples where interdomains interactions within a single polypeptide chain control allostery [38, 49]. For example, the role of UHRF1 in controlling DNA methylation requires interactions between its domains [15, 17, 19], and specifically, our previous study provided evidence for an interaction between the UHRF1-UBL and the UHRF1-SRA domain, which is required for ubiquitylation of histone H3 [9]. Studying interdomain interactions can be difficult, given the weak and transient nature of these interactions, and we thought METRIS is well-suited to measure this type of interaction. Accordingly, we tested the SRA and UBL interaction with METRIS by attaching biotinylated SRA to the substrate. For the SRA-UBL interaction, we measured an RP of 0.119*±* 0.004 for the particles, significantly higher than the 0.081 for an unmodified surface and the 0.085 we obtain with ubiquitin on the roller (Fig.4A). This represents the first direct measurement of the interaction between the SRA and UBL domains of UHRF1. We also tested a mutation to the UBL (W2V) that previous biochemical assays suggested is critical for the interaction [9, 13], and W2V had a significantly reduced the RP to 0.098*±*0.002 (Fig.4A). Fitting METRIS-K_*d*_ shows the Δ ΔG of the W2V variant is worth 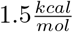 (Fig.4B and S4C) due to replacing the aromatic sidechain with the short aliphatic side chain (Fig.4D). This highlights another strength of METRIS; it is rare to assign Δ ΔG values to mutations at binding hotspots because the mutated variant binds weakly [31]. Therefore, we expect that METRIS will greatly enhance our understanding of PPIs.

### Global Fit of METRIS Analysis

We sought to generate a global fit for all of the measurements from the three independent data sets. Overall, the log-log fit of the data remained linear *(R*^*2*^ *= 0*.89) (Fig.5A), and even using this global fit, we observe agreement between fold changes and Δ ΔG within a given set of PPIs (Fig.5B). However, the METRIS-K_*d*_ values were less accurate than with the individual fitting and we could not discriminate between similar strength binders in different sets of PPIs (e.g., between DIDO1 and ORC1). These results indicating that we cannot directly compare RP values obtained for different types of PPIs and that there is likely some structural difference in each system that is not yet accounted for. However, given that each set of values had similar systematic deviations from the experimentally determined values, which is why the ΔΔG remained accurate, we realized we could apply a simple scaling factor to the METRIS-K_*d*_ values to obtain measurements that matched the experimentally determined K_*d*_. To determine the scaling factors for each interaction, we divided the published value against the METRIS-K_*d*_, and averaged them, and then multiplied the METRIS-K_*d*_ by the scaling factor and could reproduce the literature values (Fig.5B). This provides a simple way to scale METRIS-K_*d*_ values to any experimentally determined K_*d*_ values.

**Figure 5.**
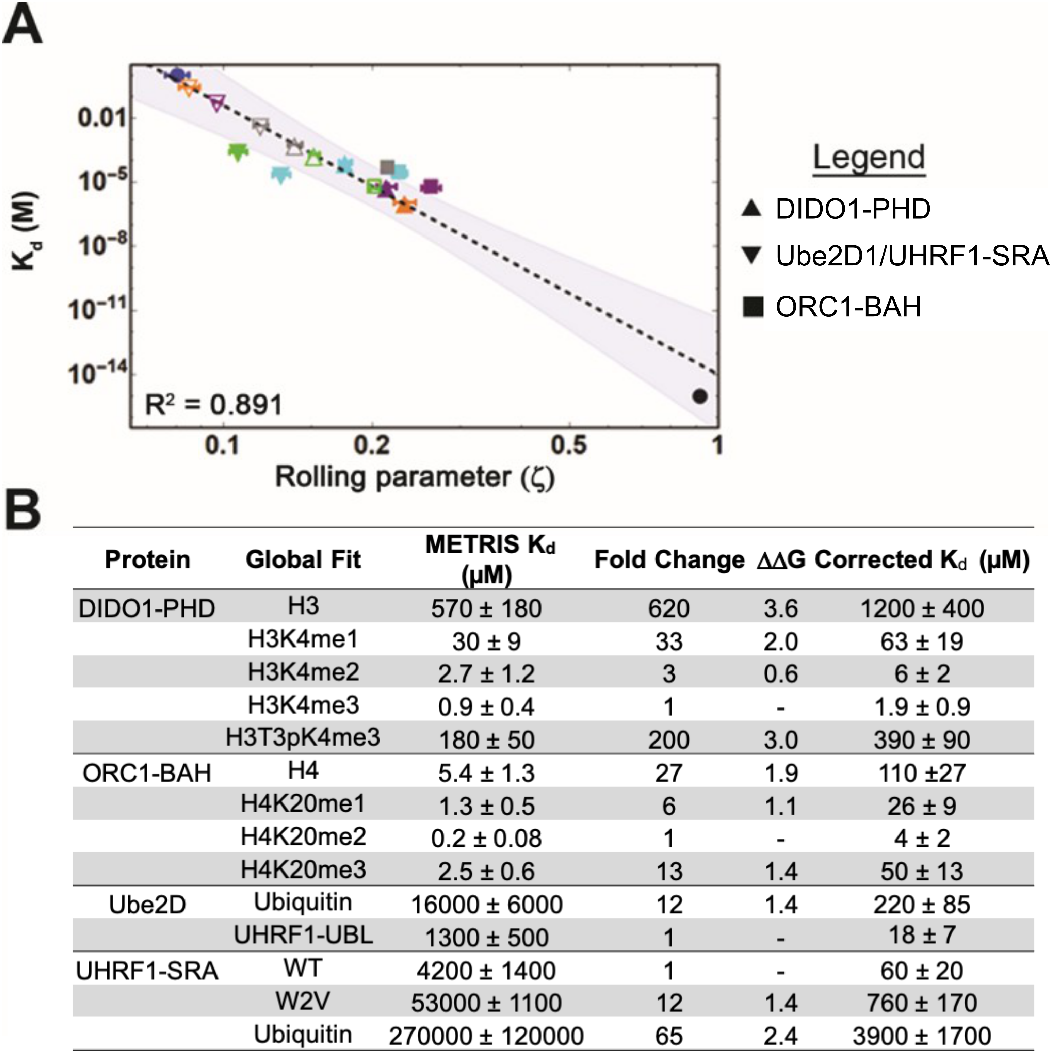
Global fit of binding partners for all METRIS experiments performed. A) Global Log-Log plot showing linearity between rolling parameter and dissociation constant for all interactions measured. B) Table of binding constants of tested interaction partners when determined from the global fit. Fold change and ΔΔ*G* are calculated in the same way as the previous example. Scaling factors are calculated by averaging the fold difference between METRIS-K_*d*_ and the published K_*d*_ for all interactions of the same type. Then the METRIS-K_*d*_ is multiplied by the scaling factor yielding the corrected K_*d*_s

## Discussion

### METRIS Can Provide New Insight into Biological Interactions

METRIS, which measures the effective mechanical friction induced by PPIs, is fundamentally different than current methodologies. This novel approach is advantageous in measuring weak interactions because it uses a very low concentration of proteins while maintaining high precision. These characteristics allow for the characterization of a vast array of PPIs, many of which were previously unmeasurable. It is difficult to overstate how transformative this will be for the study of PPIs. In this study, we demonstrate how METRIS can contribute to the study of epigenetics, by allowing us to assign ΔΔ Gs for PTMs individually and in combination, including a phospho/methyl switch is DIDO1. These values are significant because they provide a quantitative measure for the interplay between concurrent PTMs, a central premise of the epigenetic code [36].

Another area where better characterization of weak interactions will contribute significantly to understanding is in studying interdomain interactions. These types of interactions can be difficult to quantify without very resource-intensive processes, and limitations with the proteins themselves (yield or solubility) may make these interactions unmeasurable. Currently, pulldown assays, chemical crosslinking, and proximity ligation are qualitative, rarely produce quantitative data, and require mass spectrometry. Here we have measured a direct interaction between the UHRF1-UBL and SRA domains that we estimate to have a K_*d*_ 60*µ*M (Fig.5B), however, the biological context for this interaction is between two tethered domains, so an absolute value is only partially relevant. More generally, we show that METRIS can be used to measure ΔΔG for hotspot mutations, which to our understanding, could previously only be measured indirectly using high-throughput selection strategies [31]. Thus, METRIS will provide additional new data to the field of protein biochemistry and could aid in the parametrization of computational binding score functions.

An essential advantage of METRIS is resolution, precision, and sensitivity, which allows for the differentiation of ΔΔG values as small as 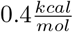. Several factors likely contribute to this robustness: 1) the rolling parameter is not inherently dependent on the protein concentration, so long as the rollers and surface are saturated. 2) The measurements have high statistical power (*≈*25 particles each with 38 RP measurements) and very low percentage error. 3) Multiple interactions between the bead and the surface amplify the friction, which may be necessary for weak interactions, and likely limits the impacts of inactive proteins on the roller and substrate. However, METRIS does have limitations, such as the reliance on literature values for extrapolating and scaling the METRIS-K_*d*_. We envision with future development, we will derive a better mathematical model that describes the relationship between protein affinity and rolling parameter as many factors will contribute to the friction, such as the number of interactions per bead or the size of the protein interaction. Despite these limitations, METRIS will be of great use to researchers studying PPIs and will provide novel information about PPIs that were previously unmeasurable.

## Acknowledgments

This work was supported by NIH grant GM126900 to B.D.S., an ACS Postdoctoral Fellowship (PF-19-027-01–DMC) to C.J.P. University of the Pacific startup funds to J.P.S. and J.S.H and an undergraduate research grant to K.R.

## Supporting Information

**Figure S1.**
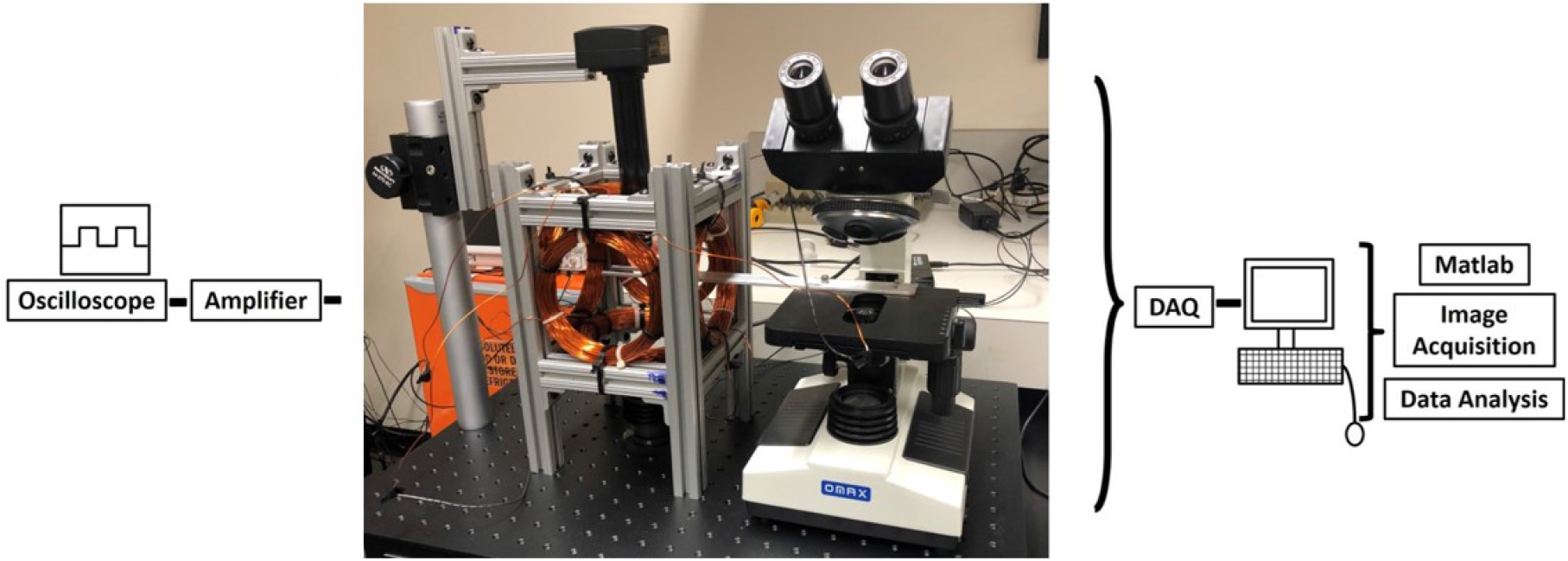
METRIS apparatus.. A) Three pairs of Helmholtz coils were mounted on an aluminum T-slot assembly. Two sinusoidal signals are generated in Matlab, passed through a DAQ, amplifier, and then to the Helmholtz coils. Visualization is accomplished using a lens tube, 10X objective, and CCD camera. A 10 mT field was utilized.

**Figure S2.**
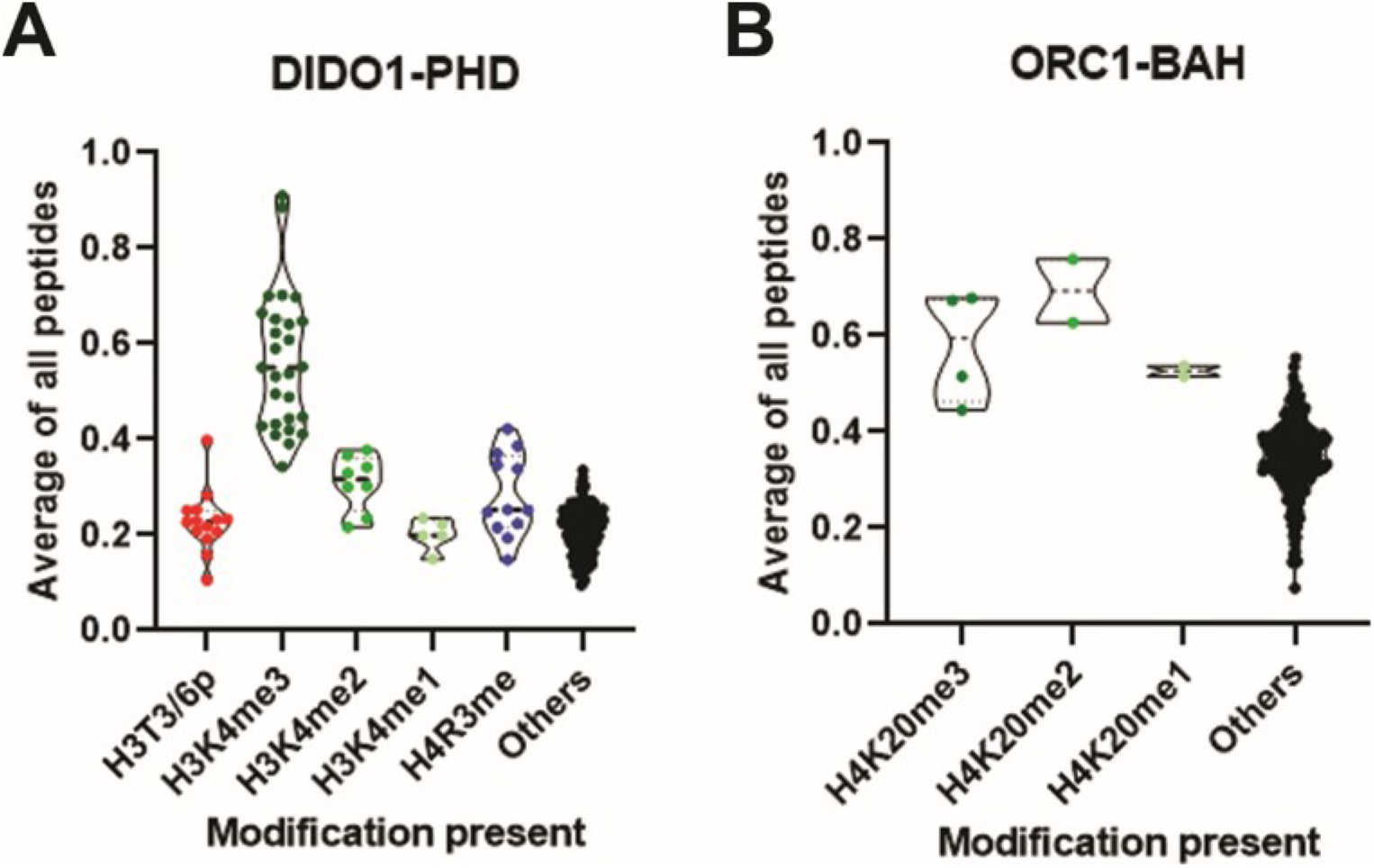
Histone peptide microarray results for all peptides for each of the chromatin reader domains.. A) and B) Normalized array signal intensities for the DIDO1-PHD (A) or ORC1-BAH (B) reader domain for peptides with the indicated modification state, with the “others” group including all other peptides. Each point represents the average value for an individual peptide. See Supplemental Data File for list of average and standard deviation for each peptide.

**Figure S3.**
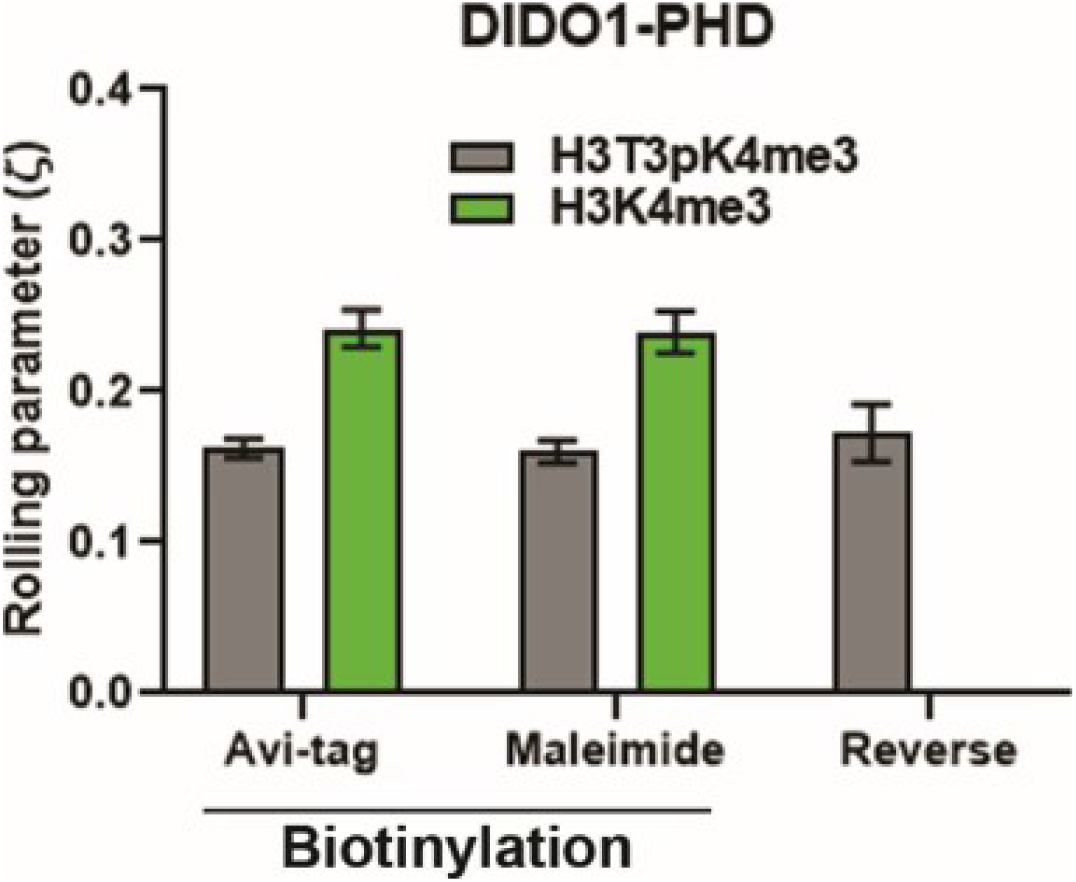
Results for tests using alternative biotinylation or functionalization strategies using the DIDO1-PHD reader module.. Results for METRIS measurements taken for H3K4me3 and H3T3pK4me3 interactions with DIDO1-PHD using different experimental designs for either biotinylation of avi-tagged GST-PHD (left two bars) or biotin-maleimide induced labeling of GST-PHD constructs. The last bar (reverse) is the result for when the functionalization of the bead and slide is reversed (peptide on the substrate and DIDO1-PHD on the roller).

**Figure S4.**
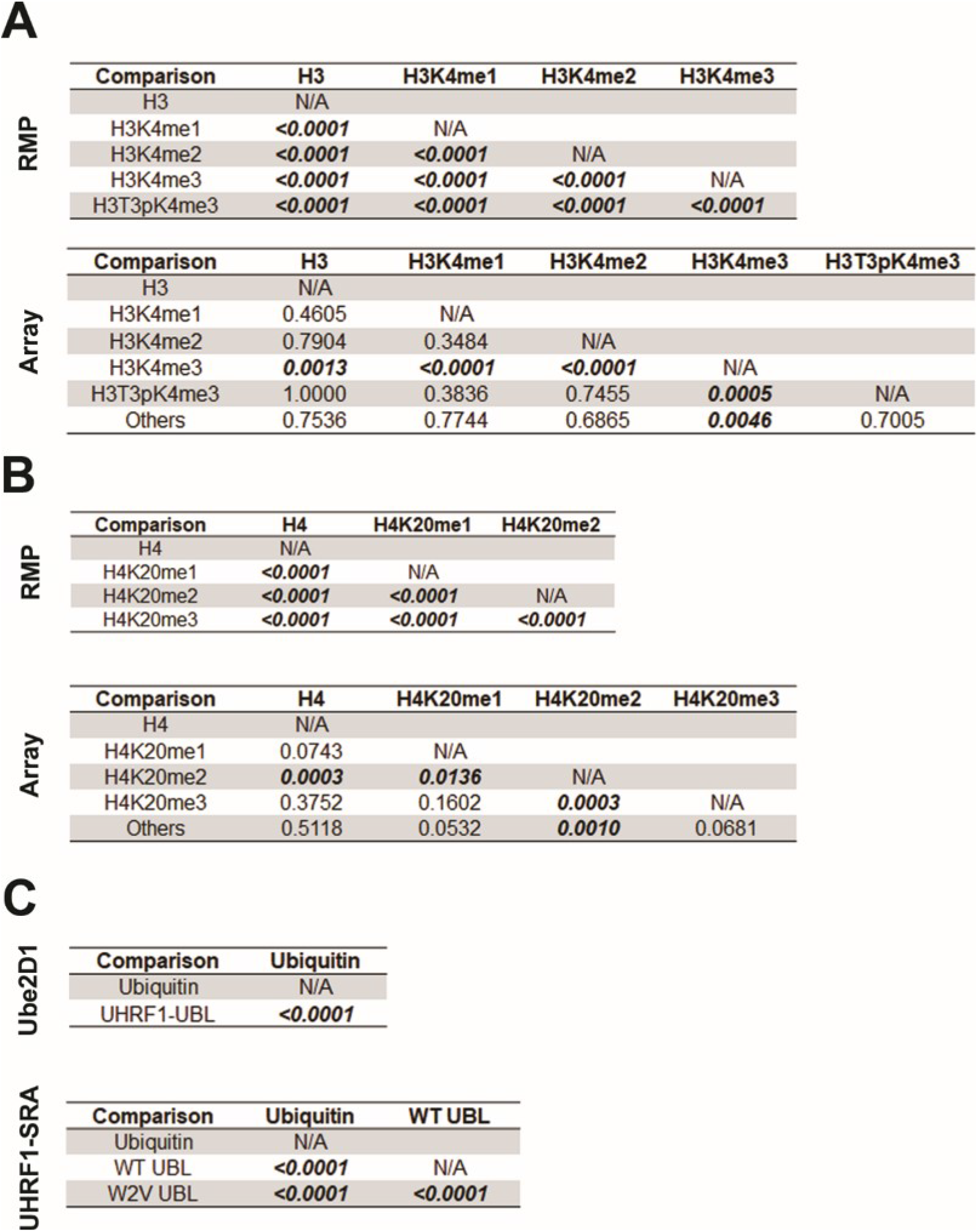
Statistical analysis of results from METRIS measurements and histone peptide microarray results.. Results from statistical tests comparing the indicated pairs of roll parameters (METRIS tables and tables labeled Ube2D1 and UHRF1-UBL) or microarray results (array tables). A Student’s T-test (unpaired, two-tailed) was used to derive the shown p-values.

